# Dermatomycosis associated with *Nannizziopsis arthrosporioides* in a breeding colony of gecko (*Correlophus ciliatus* and *Rhacodactylus auriculatus*)

**DOI:** 10.64898/2026.02.12.705528

**Authors:** Junpei Nagao, Tsuyoshi Hosoya, Kyung-Ok Nam, Genki Ishiyama, Sho Kadekaru, Yumi Une

**Author notes:** Corresponding Author: Yumi Une, DVM, PhD., The Animal Disease Research and Support Association, 2–1–2 Den-enchofu, Oota, Tokyo 145–0071, Japan.

## Abstract

This report describes lethal Nannizziopsis-associated dermatomycosis in a breeding colony of the family Diplodactylidae (*Correlophus ciliatus* and *Rhacodactylus auriculatus*). After introducing one gecko from overseas, three with indirect contact history died due to severe skin lesions. Extensive lesions were observed on the toe pads and ventral surface, along with necrotic dermatitis and cellulitis associated with fungi forming hyphae. Subsequently, four geckos developed diarrhea, melena, emaciation, and fungal dermatitis of the toe pads and died. Histopathologically, the fungal morphologies observed in the skin lesions of the seven geckos were consistent, and *Nannizziopsis arthrosporioides* was isolated and identified in two of them. To our knowledge, this is the first report of a fatal outbreak of *N. arthrosporioides* in geckos.

---

Recently, fungal infections in wild and captive reptiles have gained significant attention [13]. This is because emerging fungal diseases, such as ophidiomycosis (snake fungal disease), are increasingly being recognized as threats to ecosystems and biodiversity [13]. In veterinary clinical practice, these infectious diseases are frequently encountered, often causing severe skin lesions and frequently leading to death [2, 8]. Furthermore, fungi are opportunistic pathogens, requiring consideration of host factors in disease onset and severity, and effective treatments are not always established [2, 8]. In recent years, a fungal group with high affinity for skin keratin, previously known as *Chrysosporium* anamorph of *Nannizziopsis vriesii* (CANV), has gained attention as a cause of dermatomycosis in reptiles. In 2013, Sigler et al. molecularly classified CANV into the following three distinct fungal genera: *Nannizziopsis, Paranannizziopsis*, and *Ophidiomyces* [15, 16]. Furthermore, these fungal genera are known to possess distinct host ranges and pathogenicity. *Nannizziopsis* is primarily known as a fungus that affects lizards; however, it has now been detected in snakes, turtles, and many other types of reptiles [16]. The genus *Nannizziopsis* comprises at least 12 species, but the effects of each species on specific reptiles remain unknown [3, 15, 16]. Particularly, *N. arthrosporioides* has been reported only in five reptile species, and its infection status and pathogenicity remain poorly understood [3-5, 10, 16]. Therefore, in this study, we conducted pathological and microbiological examinations of deceased geckos to clarify the pathological findings and pathogenesis of *N. arthrosporioides* infection in two species of geckos belonging to the family Diplodactylidae.

The outbreak was confirmed at a breeding facility housing approximately 30 *Correlophus ciliatus* and *Rhacodactylus auriculatus*. First, a captive-bred male *C. ciliatus* (Case 1), imported from overseas at the end of May 2024, developed severe abdominal skin disease in early June and died. One month later, in July, two geckos (Cases 2–3) that had indirect contact with Case 1 successively developed skin lesions similar to those in case 1 and died following an acute course. Approximately 5 months after Case 1 died, four geckos with an indirect contact history with Case 1 developed diarrhea, melena, emaciation, and dermatitis on their toe pads. They became increasingly weak and eventually died one after another. The two species of geckos, totaling seven geckos, exhibited differences in the spread of skin lesions; however, all were histopathologically diagnosed with necrotic dermatitis with homogeneous characteristics. No changes were made to the husbandry conditions, such as diet or environment, before the outbreak. Additionally, the geckos were maintained in a breeding room at a temperature of 26–28 °C and a humidity of 40–50%. All geckos were housed in individual breeding boxes with pet sheets as substrates. They were fed a specialized artificial diet for *C. ciliatus*. These are standard husbandry practices for the family Diplodactylidae, and no issues were encountered during their care. The six geckos that developed skin lesions (Cases 2–7) and the newly imported gecko (Case 1) had indirect contact through shared access to the same enclosure and feeding spoons. No skin lesions were observed in other geckos that had no contact with the newly introduced gecko.

Geckos (two *R. auriculatus* and five *C. ciliatus*) were subjected to pathological examination. The geckos were adults aged 18–36 months, comprising four males and three females (Table 1). Cases 6 and 7 were treated with antibiotics and antifungal drugs, but their response to treatment was poor. These seven geckos were completely dissected.

**Table 1.**
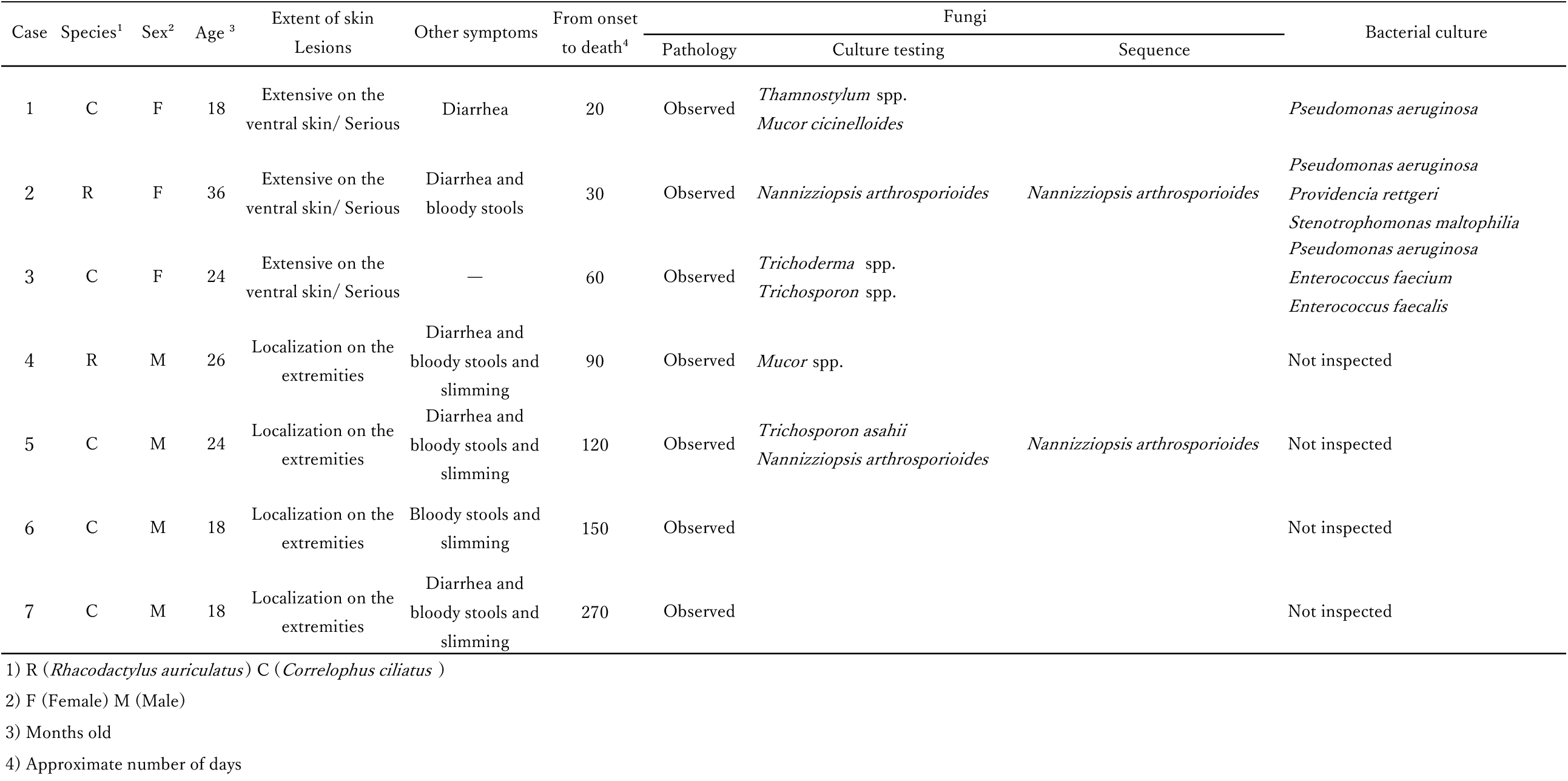
Profile and pathogen testing of cases.

For histopathological examination, the entire body, including the skin and major organs, was fixed in 10% neutral-buffered formalin. Following standard procedures, they were embedded in paraffin, thinly sectioned, and stained with hematoxylin and eosin to prepare histopathological specimens. Specific staining (Grocott staining) was performed as required. Five cases with skin lesions underwent mycological examinations (isolation culture, morphological observation and identification, and genetic analysis). Three cases underwent bacteriological examinations (isolation culture and mass spectrometry).

Three geckos (Cases 1–3) exhibited extensive, severe skin lesions with map-like discoloration of the entire abdominal trunk, showing dark red or blackish-green hues, and thickened, coarser skin. The subcutaneous tissue at the site of the skin lesion appeared red (cellulitis). Additionally, the toe pads on the limbs were dark red and hard with no subdigital lamellae, resulting in a plate-like appearance. Skin discoloration was also observed around the cloaca and ventral skin of the tail in some geckos (Figure 1). Small lesions accompanied by hemorrhagic halos approximately 2 mm in diameter were also observed. In Case 2, fibrin deposition within the body cavity, adhesion between organs, hepatic degeneration, pinhead-sized white nodules, and localized duodenal hemorrhage were also noted. Four geckos (Cases 4–7) exhibited toe pad discoloration on all limbs with loss of subdigital lamellae, were severely emaciated (Figure 1), and had small, localized trunk lesions; severe necrotic enteritis was observed in Case 6.

**Figure 1.**
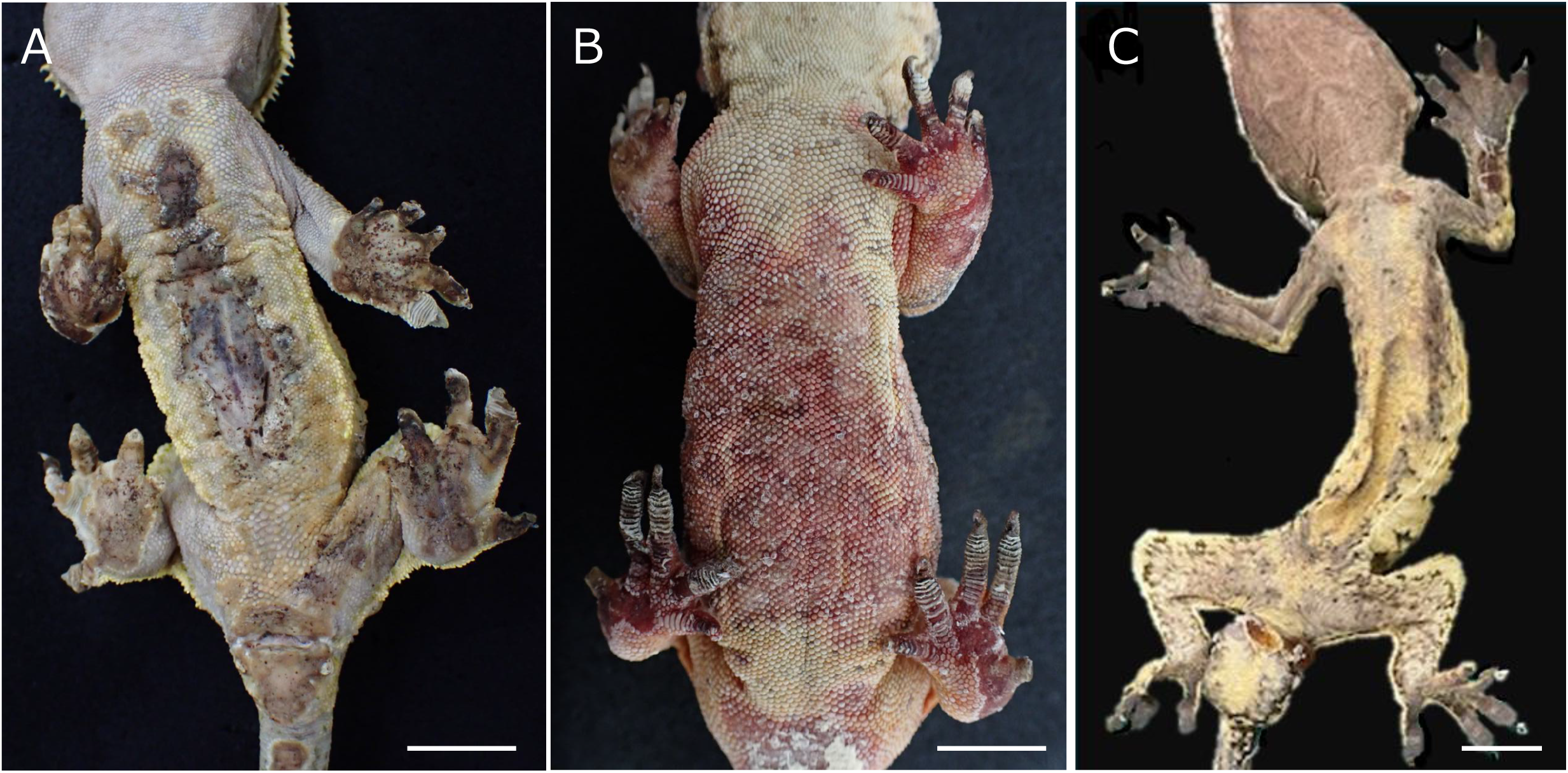
Appearance of the geckos. Skin lesions on the ventral surface of the trunk and toe pads. A: Crested gecko (Case 1)–imported animal, where symptoms were first observed. Skin lesions were observed on the abdominal skin of the trunk and the webbing between the toe pads of the limbs, and spread to the skin surrounding the cloaca and tail. The skin surrounding the ulcer is thickened. B: The skin lesions in Case 2 were red, with severe necrosis and phlegmon. C: Severe emaciation with discoloration of the toe pads on the limbs and loss of the subdigital plates. (Case 7), All scale bar:10 mm

Although the severity of the skin lesions varied, similar histological features were observed. The epidermal layer of the affected skin was completely absent, revealing layered fungal proliferation with hyphae frequently extending into the dermis and subcutaneous tissue. Necrosis of the skin and subcutaneous tissue was extensive; however, inflammatory cell infiltration was mild. Fungi exhibited relatively uniform hyphal widths and septa, with right-angled or obtuse branching (Figure 2). Bacterial proliferation was also observed in the lesions, particularly in Cases 1–3, where bacterial masses of various morphologies were observed at multiple locations. In Case 2, hyphae exhibiting similar morphology as those observed in the skin lesions were also found on the pancreatic serosa, liver parenchyma, and liver capsule surface, among other visceral organs. Four of the seven geckos exhibited diarrhea and melena. Among these, Cases 6 and 7 had bacterial necrotizing enteritis, but no fungi were detected in the affected areas. A detailed intestinal examination in Cases 1–5 was not possible because of postmortem changes and freezing effects.

**Figure 2.**
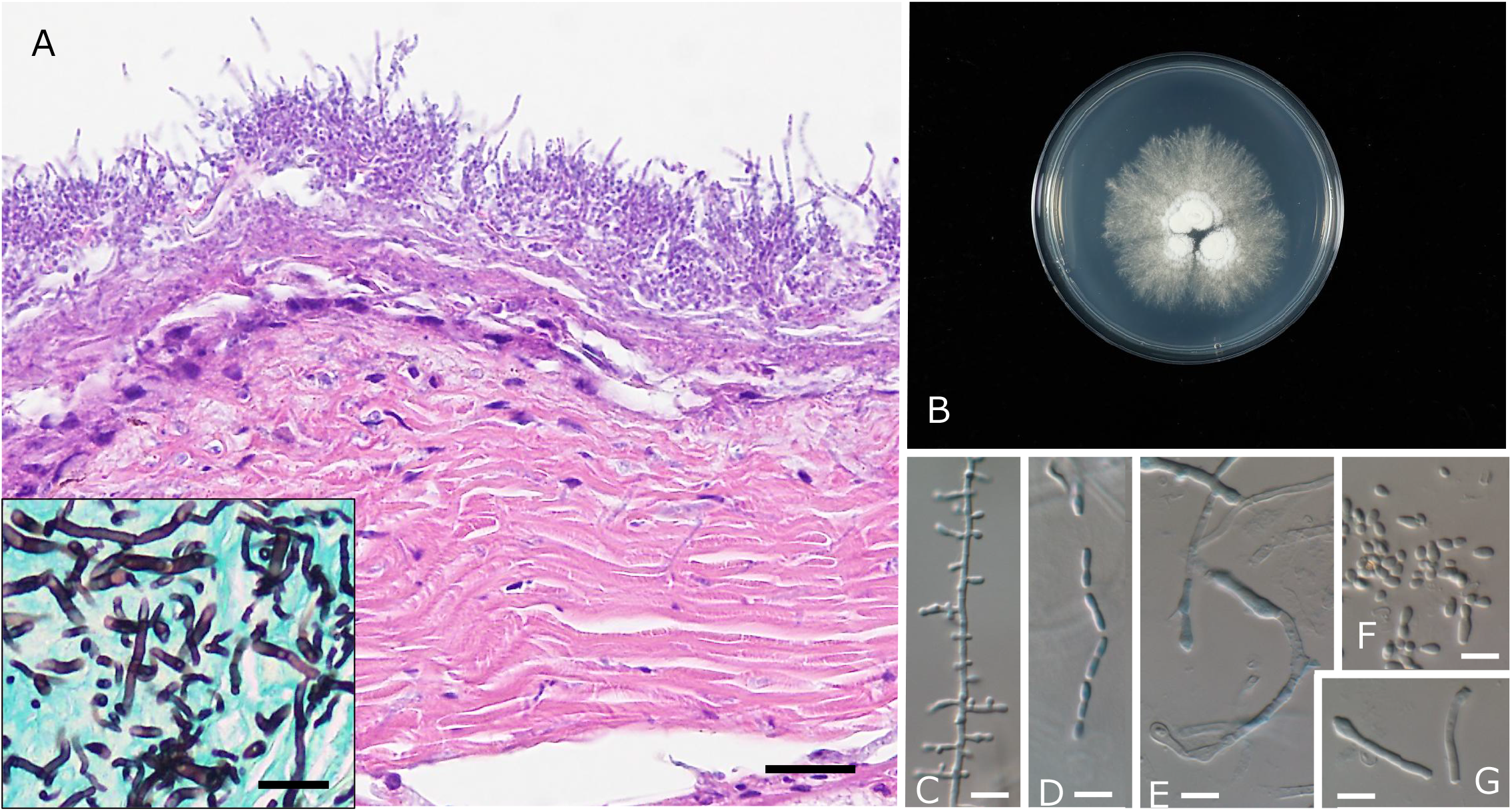
Histopathological findings of gecko skin and fungal morphology of *Nannizziopsis arthrosporioides*. A: The epidermis at the lesion site is completely absent, revealing extensive fungal proliferation. Hyphae infiltrated the subcutaneous tissue. Hematoxylin and eosin staining. Case 1, Scale bar: 100 μm. The inset shows the hyphae with septa. Branching occurred at right or obtuse angles. Grocott staining. Scale bar: 25 μm. B: Colony on potato dextrose agar. C: Typical conidiogenesis observed as a lateral protrusion. D: Arthric conidia formed by hyphae fragmentation. E: Longer arthric conidia that are produced intercalary or terminally, characterizing *N. arthrosporioides*. F: Detached conidia. Note some short clavate conidia with truncate base. G: Detached longer arthric conidia that characterize *N. arthrosporioides*. Scale bar: 10 μm.

For fungal DNA extraction, a portion of the colony with conidia formed on a potato dextrose agar (PDA; Shimadzu, Tokyo, Japan) slant was collected in a 2 mL round-bottom plastic tube and lysed using a Qiagen Tissuelyser (Hilden, Germany) using zirconia beads. DNA was extracted by incubation in 800 µL of hexadecyltrimethylammonium bromide buffer (CTAB buffer: 2% CTAB, 100 mM Tris pH 8.0, 20 mM ethylenediaminetetraacetic acid [EDTA], 1.4 M sodium chloride) at 65 °C for 60 min, and proteins were removed using a chloroform/isoamyl alcohol mixture (24:1) and centrifuged at 12,000 rpm for 15 min. The supernatant was further treated with 1,000 µL of 6 M sodium iodine buffer (NaI buffer: 6 M NaI, 50 mM Tris pH 8.0, 10 mM EDTA, 0.1 M sodium sulfite) and glass milk and stirred by a rotator for >1 hr and spun down. The precipitate was washed with 1,000 µL of ethanol/buffer solution (80% ethanol, 10 mM Tris pH 8.0, 1 mM EDTA), and finally, DNA was eluted in 120 µL of Tris-EDTA at pH 8.0 (Tris-EDTA buffer, Wako, Tokyo, Japan). The extracted DNA samples were preserved at the Center for Molecular Biodiversity Research at the National Museum of Nature and Science and are available for collaborative studies. Polymerase chain reaction (PCR) was conducted to amplify the internal transcribed spacers-5.8S ribosomal regions (ITS-5.8S rDNA) with ITS5 and ITS4 [20], and the partial actin gene with ACT1 and ACT4R primer pairs [18]. The PCR cocktail contained the following reagents: 1.0 µL of extracted DNA, 3.5 µL of DNA-free water, 5.0 µL of EmeraldAmp PCR MasterMix (Takara Bio, Kusatsu, Japan), 0.25 µL of 10 nM forward primer, and 0.25 µL reverse primer. The following protocol was applied: initial denaturation at 94 °C for 3 min; 35 cycles of 94 °C for 35 sec, 51 °C for 30 sec, and 72 °C for 60 sec; final extension at 72 °C for 10 min. The amplified PCR products were purified using ExoSAP-IT (Thermo Fisher Scientific, Waltham, MA, USA) following the manufacturer’s protocol. Sequencing reactions were carried out using an ABI PRISM 3500xl Genetic Analyzer (Applied Biosystems, Norwalk, CT, USA). The obtained sequences were assembled using the ATGC software, version 7.0.3 (Genetyx, Tokyo, Japan), and deposited in the International Nucleotide Sequence Database Collaboration.

Multiple fungal isolates were obtained from the skin of the five animals (Table 1), with isolates morphologically consistent with *Nannizziopsis* identified specifically from Cases 2 (FC-7673) and 5 (FC-7731). The strains were identical in terms of colony formation and micromorphology.

The isolate FC-7731 exhibited the following mycological features: growth was relatively slow, reaching a colony of diameter of 5–6 cm after 1 month at 25 °C; the colony appeared powdery due to abundant conidial production and was white with a yellowish tinge. Conidia were thallic, single-celled, and borne singly or in chains, either terminally or intercalary; they were typically ellipsoid or short clavate with a truncate base, (3–)5–6 x (2–)3–4 μm, produced as lateral protrusions from hyphae leaving scars when detached. Conidia were also produced by hyphae fragmentation, forming cylindrical arthroconidia 15–28 μm long and 3–5 μm wide (Figure 2). These features are consistent with the descriptions of *N. arthrosporioides* by Stchigel et al. [16].

The sequences of ITS and actin obtained from FC-7673 and 7731 were identical where comparable. In a BLAST search of the National Center for Biotechnology Information database (https://blast.ncbi.nlm.nih.gov/Blast.cgi), the ITS (LC916016 from FC-7731 and LC916017 from FC-7673) and actin sequences (LC916018 from FC-7731 andLC916019 from FC-7373) showed 98.94%, 99.98%, 99.62, 99.59% identity with the sequence from the type material of *N. arthrosporioides* UTHSC R4263 (accession numbers NR111523 for ITS and HF547885 for actin), respectively. The actin sequence also showed more than 98% identity to other sequences of *N. arthrosporioides* (such as OR240086, PV551206, and MN182618). Based on these analyses, we identified FC-7731 and FC-7673 as *N. arthrospoirioides*. The isolates will be deposited in the Biological Resource Center, part of the National Institute of Technology and Evaluation, Japan.

Pathological tissue specimens from all seven skin lesions showed growth of various bacteria. Bacteria were collected from the skin of Cases 1–3, which exhibited particularly severe skin symptoms, and were submitted for culture and identification testing. Mixed infections involving bacteria such as *Pseudomonas aeruginosa, Providencia rettgeri, Stenotrophomonas maltophilia*, and *Enterococcus faecalis* were identified.

At a breeding facility, an outbreak of fungal necrotic dermatitis occurred in seven geckos over a specific period following the introduction of a new gecko. Histopathologically, identical filamentous fungi were observed at the skin lesion sites. *N. arthrosporioides* was isolated and mycologically identified. Therefore, this case was diagnosed as an epidemic *N. arthrosporioides* infection in geckos. This infectious disease has been reported in turtles, snakes, lizards, and humans; however, this is the first confirmed case in geckos [3-6, 10, 16]. The causative fungi of dermatomycoses, previously classified as CANV, are believed to be transmitted through contact with infectious spores. This contact occurs either directly during reproductive activities such as mating or indirectly via contaminated objects, with the latter often serving as the primary source of infection. [10, 12, 14, 16,17]. In this case, all geckos were kept separately. However, as the disease manifested only in the newly imported animal and those sharing feeding practices and housing equipment among approximately 30 geckos, it was inferred that this outbreak resulted from horizontal transmission via indirect contact. Outbreaks of *Nannizziopsis* fungi in reptiles within the same facility have been reported in multiple species, including bearded dragons, green iguanas, crocodiles, and leopard geckos [9, 14, 15, 17]. These species include *N. guarroi, N. vriesii, N. crocodile*, and *N. chlamydospora*, which have been reported to cause fatal epidemics. Particularly, infection with *N. guarroi* is highly pathogenic in bearded dragons and green iguanas [7], and it is suggested that *N. arthrosporioides*, as confirmed in this study, may exhibit similar pathogenicity. Therefore, *N. arthrosporioides* requires careful attention, and the establishment of rapid diagnostic methods and treatments is urgently needed. Regarding the pathogenesis of this case series, for Cases 1–3, we surmised that death resulted from tissue damage due to extensive and severe skin lesions, combined with deep and disseminated fungal infections or sepsis caused by bacterial infections. In cases 4–7, although necrotic dermatitis involving the toe pads was limited, local skin damage was severe, accompanied by significant bacterial infection, and the animals’ nutritional status was extremely poor. Additionally, bacterial necrotic enteritis, which is considered a cause of bloody diarrhea, was observed. These factors were judged to be strongly involved in the causes of death. All bacteria isolated from the lesion site were environmental and not those indicative of an epidemic. Therefore, bacterial infections were presumed to be secondary opportunistic infections following dermatitis caused by *N. arthrosporioides*. Information on *N. arthrosporioides* is extremely scarce, but previous reports indicate that it primarily causes skin lesions, such as ulcers, with cases of deep fungal infections also documented [3-5, 10, 16]. In this case, invasive proliferation of the dermis and deep skin layers was frequently observed, and severe deep fungal infections were noted. The extent of fungal proliferation varies between cases, but *N. arthrosporioides* itself, similar to other *Nannizziopsis* species, can cause death as a deep-seated fungal infection [1]. They are also thought to destroy skin defense mechanisms, providing opportunities for other microorganisms to invade [17]. Infections caused by fungi of the genus *Nannizziopsis* are reportedly triggered by factors such as inappropriate diet, housing conditions, and environmental stress. However, in this study, a sudden outbreak occurred in a breeding facility that maintained stable reproduction for a long period. This led us to conclude that *N. arthrosporioides* possesses high infectivity and pathogenicity [1, 12, 17]. CANV is a novel infectious disease that affects not only reptiles but also immunocompromised humans [2, 11, 16]. To date, *N. arthrosporioides* has been considered an infectious disease that affects reptiles. However, recent reports of human infections suggest a broad host range for *N. arthrosporioides* [6], necessitating caution as a zoonotic disease [6, 10]. Reptiles, including members of the family Diplodactylidae, are popular pets actively traded internationally [19]. This international trade in reptiles may represent a new transmission route for zoonotic fungal diseases, necessitating public health risk assessments [3].

## ACKNOWLEDGMENTS

This work was supported by the Health and Labour Science Research Grants (21HA2001).

## DATA AND MATERIALS AVAILABILITY

All data for this study are described in the main text of the paper.

## ETHICS STATEMENT

This paper was prepared with the consent of all co-authors, and there are no ethical or privacy restrictions.

## REFERENCES

1. Bowman, M. R., Pare, J. A., Sigler, L., Naeser, J. P., Sladky, K. K., Hanley, C. S., Helmer, P., Phillips, L. A., Brower, A., Porter, R. 2007. Deep fungal dermatitis in three inland bearded dragons (Pogona vitticeps) caused by the Chrysosporium anamorph of Nannizziopsis vriesii. Med Mycol 45: 371–376.

2. Cabanes, F. J., Sutton, D. A., Guarro, J. 2014. Chrysosporium-related fungi and reptile: a fatal attraction. PLoS Pathog 10: e1004367.

3. Chen, Y. H., Chi, M. J., Sun, P. L., Yu, P. H., Liu, C. H., Cano, L., Jose, F., Li, W. T. 2020. Histopathology, Molecular Identification and Antifungal Susceptibility Testing of Nannizziopsis arthrosporioides from a Captive Cuban Rock Iguana (Cyclura nubila). Mycopathologia 185: 1005–1012.

4. Christman, J. E., Alexander, A. B., Donnelly, K. A., Ossiboff, R. J., Stacy, N. I., Richardson, R. L., Case, J. B., Childress, A. L., Wellehan, J. F. 2020. Clinical manifestation and molecular characterization of a novel member of the Nannizziopsiaceae in a pulmonary granuloma from a Galapagos tortoise (Chelonoidis nigra). Front Vet Sci 7: doi: 10.3389/fvets.2020.00024.

5. Durante, K., Sheldon, J. D., Adamovicz, L. A., Roady, P. J., Keller, K. A. 2023. Systemic Nannizziopsis arthrosporioides in an African side-neck turtle (Pelomedusa subrufa) J. Herpetol. Med. Surg 33: 223–228.

6. Ende, B. G., Rodrigues, A. M., Hahn, R. C., Hagen, F. 2023. A surprising finding: the curious case of a tongue lesion misdiagnosed as paracoccidioidomycosis. Rev Iberoam Micol 40: 10–14.

7. Gentry, S. L., Lorch, J. M., Lankton, J. S., Pringle, A. 2021. Koch’s postulates: Confirming Nannizziopsis guarroi as the cause of yellow fungal disease in Pogona vitticeps. Mycologia 113: 1253–1263.

8. Hatt, J. M. 2010. Dermatological diseases in reptiles. Schweiz Arch Tierheilkd 152, 123–130.

9. Hill, A. G., Sandy, J. R., Begg, A. 2019. Mycotic dermatitis in juvenile freshwater crocodiles (Crocodylus johnstoni) caused by Nannizziopsis crocodile. J Zoo Wildl Med 50: 225–230.

10. Keller, K. A., Adamovicz, L., Johnson-Delaney, C., Terio K A. 2025. Nannizziopsis arthrosporioides infection mimicking ophidiomycosis in ball pythons (Python regius). Med Mycol Case Rep 50: doi: 10.1016/j.mmcr.2025.100733.

11. Nourrisson, C., Roux, V. M., Cayot, S., Jacomet, C., Bothorel, C., Pilon, L. A., Lesens, O., Poirier, P. 2018. Invasive infections caused by Nannizziopsis spp. molds in immunocompromised patients. Emerg Infect Dis 24: 549–552.

12. Pare, A., Coyle, K. A., Sigler. L., Maas, M. K., Mitchell, R. L. 2006. Pathogenicity of the Chrysosporium anamorph of Nannizziopsis vriesii for veiled chameleons (Chamaeleo calyptratus). Medical Mycology 44: 25–31.

13. Peterson, N. R., Rose, K., Shaw, S., Hyndman, T. H., Sigler, L., Kurtboke, D I., Llinas, J., Littleford-Colquhoun, B. L, Cristescu, R., Frere, C. 2021. Cross-continental emergence of Nannizziopsis barbatae disease may threaten wild Australian lizards. Sci Rep 10: 20976: doi: 10.1038/s41598-021-86463-0.

14. Schmidt-Ukaj, S., Loncaric, I., Spergser, J., Richter, B., Hochleithner, M. 2016. Dermatomycosis in three central bearded dragons (Pogona vitticeps) associated with Nannizziopsis chlamydospore. J Vet Diagn Invest 28: 319–322.

15. Sigler, L., Hambleton, S., Pare, J. A. 2013. Molecular characterization of reptile pathogens currently known as members of the Chrysosporium anamorph of Nannizziopsis vriesii complex and relationship with some human-associated isolates. Journal of Clinical Microbiology 51: 3338–3357.

16. Stchigel, A. M., Sutton, D. A., Cano-Lira, J. F., Cabanes, F. J., Abarca L., Tintelnot, K., Wickes, B. L., Garcia, D., Guarro, J. 2013. Phylogeny of Chrysosporia infecting reptiles: proposal of the new family Nannizziopsiaceae and five new species. Persoonia 31: 86–100.

17. Toplon, D. E., Terrell, S. P., Sigler, L., Jacobson, E. R. 2013. Dermatitis and cellulitis in leopard geckos (Eublepharis macularius) caused by the Chrysosporium anamorph of Nannizziopsis vriesi. Vet Pathol 50: 585–589.

18. Voigt, K., Wöstemeyer, J. 2000. Reliable amplification of actin genes facilitates deep-level phylogeny. Microbiological Research 155: 179–195.

19. Warwick, C. 2014. The morality of the reptile “pet” trade. J Anim Ethics 4:74–94.

20. White, T. J., Bruns, T., Lee, S. and Taylor, J. 1990. Amplification and direct sequencing of fungal ribosomal RNA genes for phylogenetics. pp. 315–322. In: PCR Protocols: A Guide to Methods and Applications (Innis M. A., Gelfand D. H., Sninsky J. J. and White T. J. eds.), Academic Press, San Diego.

